# TLR4 promotes liver fibrosis in metabolic dysfunction-associated steatohepatitis by a mechanism independent of hepatocytes and inflammatory cells

**DOI:** 10.1101/2025.09.25.678670

**Authors:** Eliza Elza Altvater, Daniah Kojh, Ryuan Huang, Jorge Matias Caviglia

## Abstract

In metabolic dysfunction-associated steatotic liver disease (MASLD), liver fibrosis is the most important prognostic factor, and fibrosis can lead to hepatic cirrhosis and cancer. In the liver toll-like receptor 4 (TLR4) signaling promotes fibrosis. To investigate the role of TLR4 in the development of MASLD, we used a mouse model of MASLD, and we deleted the *Tlr4* gene either in the whole body or selectively in inflammatory cells or hepatocytes. Mice with a whole-body deletion of *Tlr4* developed MASH with steatosis, hepatocellular injury and inflammation; however, *Tlr4* deletion prevented the deposition of fibrosis. *Tlr4* deletion on immune cells or in hepatocytes did not prevent fibrosis. These results suggest that activation of TLR4 in other hepatic cells, possibly hepatic stellate cells, or in a combination of cells promotes liver fibrosis in MASLD.

## INTRODUCTION

Metabolic dysfunction-associated steatotic liver disease (MASLD)^3^ is the most prevalent liver disease in the US and worldwide [1, 2]. MASLD includes metabolic dysfunction-associated steatotic liver, metabolic dysfunction-associated steatohepatitis (MASH), and cirrhosis [3, 4]. MASLD can lead to liver failure and cancer [3]. In patients with MASLD, fibrosis is the primary predictor of disease progression, morbidity, and mortality [3]. Liver cancer develops in fibrotic livers in 70-80% of the cases and the risk of developing hepatocarcinoma increases with more advanced fibrosis stages [3, 5, 6]. Importantly, collagen type I, the most abundant extracellular protein in liver fibrosis, promotes tumor development, implicating fibrosis in the pathophysiology of hepatocarcinoma [7].

Toll-like receptor 4 (TLR4) activation, as well as activation of other TLRs, promote the development of liver fibrosis [8-10]. Mice with a loss-of-function mutation in the *Tlr4* gene had reduced liver fibrosis in hepatotoxic and cholestatic models of liver injury [11]. In mouse models of MASLD, previously called non-alcoholic fatty liver disease (NAFLD), *Tlr4* inactivating mutations or deletion of *Tlr4* reduced fibrosis [12-14]. Moreover, in humans, TLR4 variants affects fibrosis risk [15, 16].

TLR4 is activated by lipopolysaccharide (LPS), a component of intestinal bacteria, whose plasma concentration increases in obesity and MASLD [16, 17]. Additionally, in obesity, expanded adipose tissue and accelerated lipolysis increase the release of fatty acids, including saturated fatty acids, thereby increasing their levels in blood [18]. Saturated fatty acids, which resemble the lipid A moiety of lipopolysaccharide, also activate TLR4 [19, 20].

Therefore, in MASLD and obesity, which often co-exist, TLR4 can be activated by obesity-related ligands and TLR4 can promote liver fibrosis. Until now, the contribution of the various hepatic cell populations to TLR4-mediated fibrosis in MASLD has been less studied. In liver, TLR4 is expressed in hepatocytes, endothelial cells, and at higher levels in macrophages and hepatic stellate cells, two cell populations that are important in inflammation and fibrogenesis [11, 21, 22].

In non-MASLD mouse models of liver fibrosis, TLR4 activation in hepatic stellate cells sensitized the cells to TGFβ, contributing to their activation, differentiation into myofibroblasts, and production of extracellular matrix [11]. In a mouse model of MASLD/NAFLD, deletion of *Tlr4* in hepatocytes reduced steatosis and hepatocellular injury; however, fibrosis was not assessed in that study [23]. In a different mouse model of MASLD/NAFLD, driven by a diet high in sucrose, saturated fat, and cholesterol, deletion of *Tlr4* in hepatocytes decreased fibrosis but other cell populations were not evaluated [24].

It has been proposed that TLR4 antagonists could be used to treat liver fibrosis. However, TLR4 plays a role in the immune response, and thus blocking TLR4 signaling may increase the risk of infections by gram-negative bacteria [25]. Therefore, it would be useful to determine in which of the liver cells TLR4 activation promotes fibrosis in MASLD.

To further investigate the role of TLR4 in the development of fibrosis in MASLD and to determine whether TLR4 in hepatocytes or immune cells promoted the development of fibrosis, we used a model of MASLD that we developed [26, 27]. This MASLD/NAFLD mouse model shows increased TLR4 expression as well as gut microbiota dysbiosis with an increase in LPS-producing gram negative bacteria; in addition, mice with MASH have increased plasma fatty acids [26, 27]. Using this model, we evaluated the role of TLR4 in fibrosis by deleting *Tlr4* either globally in all cells, or selectively in bone marrow-derived immune cells or in hepatocytes.

## METHODS

### Animals and diets

*Tlr4 null* mice (TLR4KO) were generated by Shizuo Akira [28]. TLR4 floxed mice were generated by Joel Elmquist [23]. Mice carrying the Ay mutation in a C57BL6/J background (B6.Cg-*A*^*y*^/J mice, The Jackson Laboratories 000021) and Vav-iCre mice, expressing the Cre recombinase in hematopoietic cells (B6.Cg-*Commd10*^*Tg(Vav1-icre)A2Kio*^/J, The Jackson Laboratories 008610) were obtained from The Jackson Laboratory. Adeno-associated virus (AAV) serotype 8 directing the expression of the Cre recombinase driven by the thyroglobulin promoter (AAV8-TBG-Cre) were used to selectively express the Cre recombinase in hepatocytes (Addgene 107787-AAV8). TLR4ΔHep mice were produced by administering the AAV8-TBG-Cre intravenously by retroorbital injection to *Tlr4* floxed mice. Control mice were administered a null AAV created with an empty vector (Addgene 105536-AAV8). TLR4ΔBM were generated by crossing *Tlr4* floxed mice with Vav-iCre mice; *Tlr4* floxed not carrying the Vav-iCre transgene were used as controls. We crossed these strains with Ay mice for MASH induction; littermates with genotype aa, identified by their black coat color, were used as lean controls.

MASH was induced in Ay mice by feeding them a Western-type diet (Inotiv, Teklad TD.88137) and giving them a solution containing fructose (23 g/l) and glucose (19 g/l), equivalent to high fructose corn syrup. Lean control aa mice were fed a low fat and fructose diet (Inotiv, Teklad TD.08485) and regular water. Mice were fed these diets for 16 weeks, starting at 8 weeks of age. Mice were maintained in cages with wood chip bedding, at a temperature of 20–23 °C, with a 12 h light/12 h dark cycle. All data reported correspond to male mice.

Animal care and procedures followed the Guide for the Care and Use of Laboratory Animals and were approved by the IACUC of Columbia University or Brooklyn College.

### Tissue collection

Mice were fasted with access to water for 5 hours before tissue collection. Mice were euthanized with carbon dioxide. Blood was collected from the inferior vena cava. Livers were dissected, weighed, and parts of several lobes were either fixed in formalin for histology or flash frozen in liquid nitrogen for other analyses. Fat pads were dissected and weighed.

### Evaluation of livers for MASH

Liver tissue corresponding to multiple lobes, which were fixed in formalin 10% for 24 hours, were embedded in paraffin and sections were made and mounted on slides (Columbia University Histopathology Core). Sections were stained with hematoxylin and eosin, Masson’s Trichrome, or picrosirius red [27]. Images were obtained using 4X or 10X objectives and either a QImaging Retiga or an Olympus DP28 camera.

Liver steatosis was evaluated by histopathology and by measuring liver triacylglycerols by extracting the lipids and quantifying triacylglycerols with an Infinity Triglycerides Reagent biochemical kit (ThermoScientific TR22421) [26, 29].

Liver injury was assessed by measuring plasma ALT and/or AST activity using kits BQ-Kits BQ004A and GenWay GWB-BQK284 [26].

Inflammation- and fibrosis-related gene expression was assessed by QPCR as described [26].

For collagen quantification, the picro-sirius red-stained sections were imaged using polarized light to specifically detect collagen. Collagen was quantified morphometrically as previously described [26, 27]. Collagen was also assessed by quantification of liver content of hydroxyproline as described [26, 27]

### Statistical analysis

Statistical analyses were conducted using one-way ANOVA. When variance differed significantly between experimental groups, the data was log-transformed to meet the ANOVA homoscedasticity requirement. P values smaller than 0.05 were considered statistically significant. Data presented include the mean, the standard error of the mean, and individual data.

## RESULTS

### TLR4 whole-body deletion prevented liver fibrosis

To evaluate the role of TLR4 in the development of MASH-driven liver fibrosis, we first used whole-body *Tlr4 null* (knock-out, TLR4KO) mice; wild-type (WT) mice were used as control. To induce MASH with fibrosis, we used Ay mice that were either TLR4KO or WT and fed them a high-in-fat-and-fructose diet (HFFD) consisting of a Western diet and a drinking solution containing a high fructose corn syrup equivalent [26]. TLR4KO and WT mice with genotype aa (non-hyperphagic) fed a low fat and fructose diet, which do not develop MASLD, were used as controls [26]. Deletion of the *Tlr4* gene was confirmed by the lack of *Tlr4* mRNA expression in the livers of TLR4KO MASH and control mice (Fig 1A).

**Figure 1:**
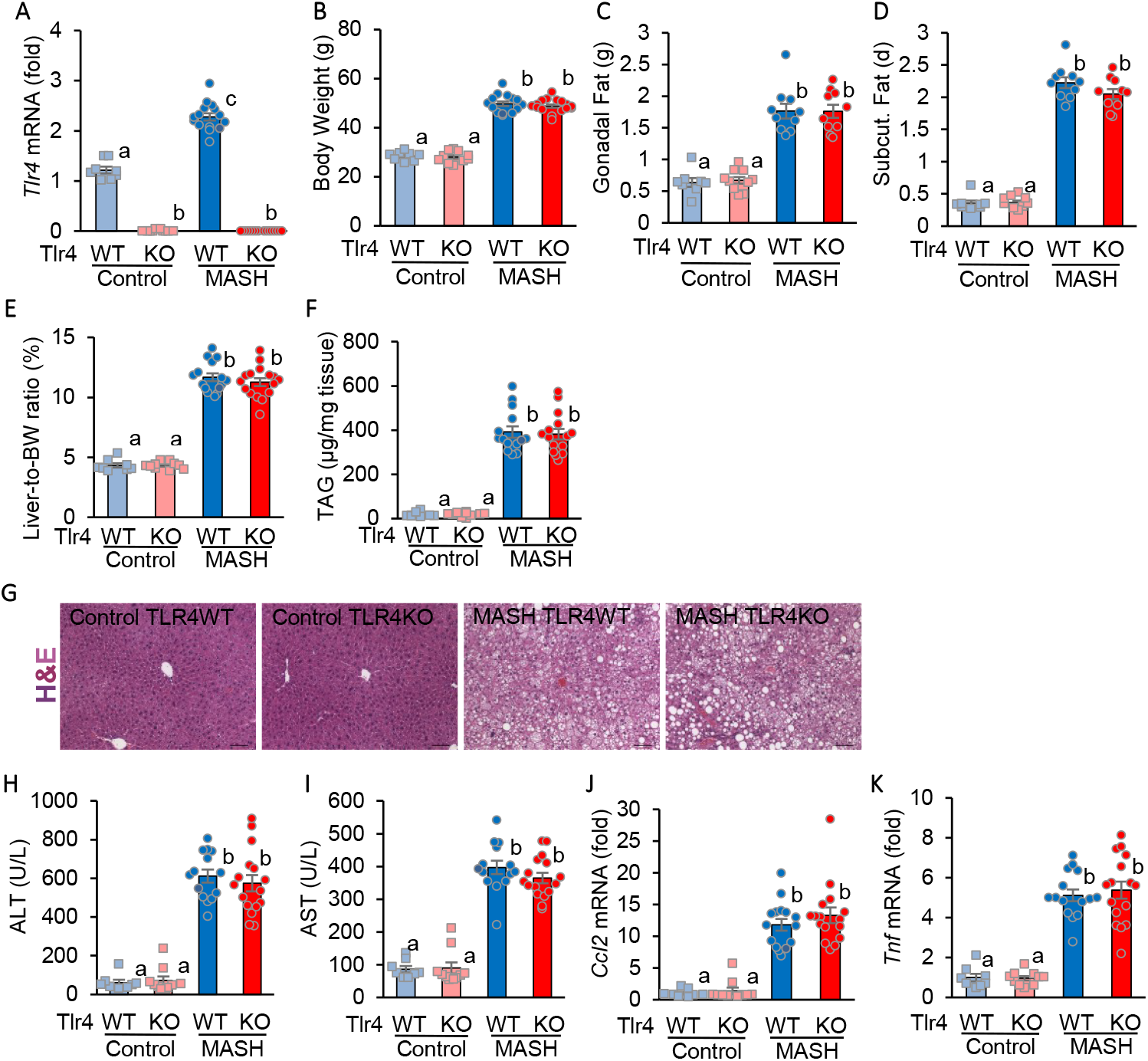
*Tlr4* deletion does not prevent obesity or metabolic dysfunction-associated steatohepatitis (MASH). *Tlr4* wild-type (TLR4 WT) and *Tlr4 null* (TLR4 KO) mice, both carrying the Ay mutation were fed a high-fat-and-fructose diet (HFFD) for 16 weeks to induce MASH; TLR4 WT or KO aa mice fed a low-fat-and-fructose diet were include as non-MASH controls (Control). (A) TLR4 deletion was verified by hepatic *Tlr4* mRNA expression. Obesity was evaluated by measuring body weight (B), and gonadal and subcutaneous fat pad weights (C;D). Hepatic steatosis was assessed by liver-to-body weight ratio (E), liver triacylglycerol (TAG) content (F), and histology (G). Hepatocellular injury was evaluated by plasma levels of AST and ALT (H; I). Inflammation was determined by hepatic mRNA expression of *Ccl2* and *Tnf* (J;K). Experimental group size were 8 (for TLR4 WT control), 10 (for TLR4 KO control), 15 (for TLR4WT MASH), and 16 (for TLR4KO MASH). Data represents data from individual mice (dots), group means (bars), and standard error of the mean (error bars). Statistical analysis was done by one-way ANOVA; different superscript letters indicate statistically significant differences with *p*<0.05.

TLR4KO MASH and WT MASH mice gained more weight and had larger gonadal fat pads than TLR4KO and WT control mice, indicating that the deletion of *Tlr4* did not prevent the development of obesity (Fig 1B-D).

TLR4KO MASH and WT MASH mice, both developed similar degrees of hepatomegaly and liver steatosis as shown by liver-to-body weight ratios, triacylglycerol content, and histology of hematoxylin and eosin-stained liver sections, as compared with lean controls (Fig 1E-G). We have previously reported that Ay mice fed the HFFD have glucose intolerance, hyperinsulinemia, and hypertriglyceridemia, meeting the criteria for diagnosis of MASLD in humans [26, 27].

TLR4KO MASH and WT MASH mice had elevated ALT and AST (Fig 1H; I), indicating that they had hepatocellular injury. The livers of the mice had inflammation, as shown by increased mRNA expression of inflammation marker genes *Ccl2* and *Tnf* (Fig 1J; K) and histology (Fig 1G). Therefore, both TLR4KO MASH and TLR4 WT MASH mice develop metabolic dysfunction-associated steatohepatitis, suggesting that TLR4 is not essential in the development of steatohepatitis.

In TLR4 WT mice, induction of MASH caused an increase in collagen deposition, as assessed by trichrome staining and measured by picro-sirius red staining of collagen and liver hydroxyproline content (Fig 2A-C). In TLR4 KO mice with MASH, deletion of *Tlr4* prevented collagen deposition (Fig 2A-C). Expression of fibrosis-related genes *Col1a1, Lox*, and *Timp1*, was increased in mice with MASH compared with lean controls; deletion of *Tlr4* in TLR4KO MASH mice decreased the expression of those genes compared with TLR4 WT MASH mice (Fig 1D-F).

**Figure 2:**
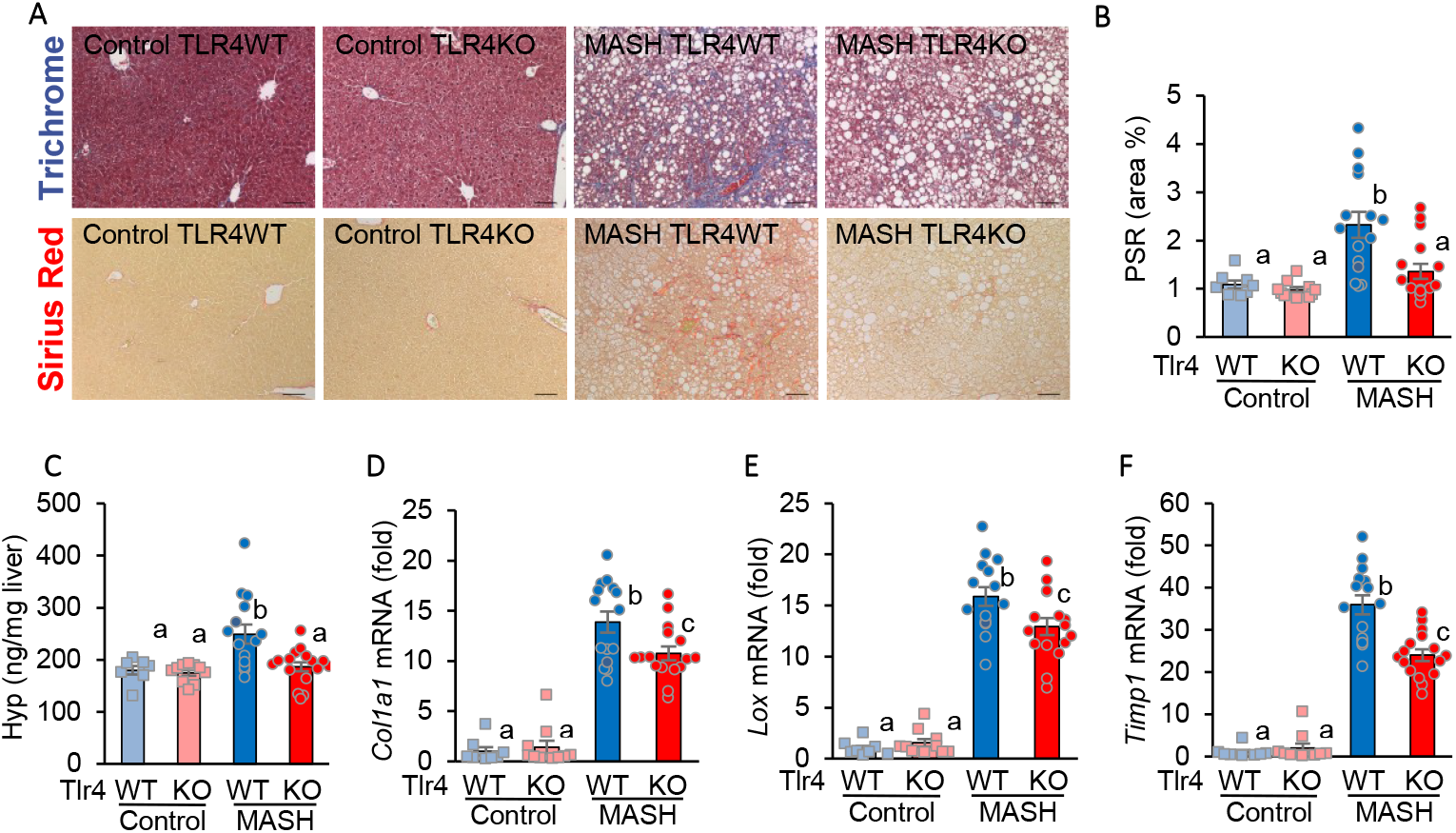
Tlr4 deletion prevents the development of liver fibrosis in MASH. TLR4 knock-out (TLR4KO) or wild-type (WT) mice were either Ay mice fed a HFFD to induce MASH, or aa fed a low-fat diet to serve as controls. Liver fibrosis was assessed by trichrome and PSR staining (A, B), hydroxyproline hepatic content (C), and mRNA expression of fibrosis-related genes *Col1a1, Lox*, and *Timp1* (D-F). N = 8/10/15/16, one-way ANOVA; different superscript letters indicate statistically significant differences with p<0.05.

Therefore, deletion of *Tlr4* prevented fibrosis in MASH, suggesting that TLR4 plays an essential role in the development of liver MASH-related liver fibrosis.

### TLR4 deletion in bone marrow-derived cells did not prevent MASH-related fibrosis

To further investigate the role of TLR4 in MASH, we evaluated the role of TLR4 in the various hepatic cell types by deleting *Tlr4* in specific cell populations and assessing the effect on the development of MASH.

To delete TRL4 in immune cells, we crossed mice that have a floxed *Tlr4* allele [23] and mice that express the Cre recombinase, directed by the Vav1 promoter, selectively in hematopoietic / bone barrow-derived cells (The Jackson Laboratory # 088610). Littermate mice carrying the floxed *Tlr4* allele but not expressing the Cre recombinase were use as controls. We designated these mice TLR4ΔBM and TLR4 ff, respectively. We generated TLR4ΔBM and TLR4 ff mice with the Ay mutation and fed them the HFFD to induce MASH; we also generated TLR4ΔBM and TLR4 ff aa mice fed the low fat and fructose diet to use as controls. We verified the *Tlr4* deletion in TLR4ΔBM by measuring its expression in spleen, which was decreased 86% compared with control TLR4 floxed mice (data not shown).

Ay mice fed the HFFD diet, both TLR4ΔBM and TLR4ff, developed obesity and MASH similarly (data not shown). Mice with MASH developed of hepatic fibrosis, which was assessed by picro-sirius red staining of liver sections and by quantification of hydroxyproline liver content (Fig 3A-C). Fibrosis was similar in TLR4ff MASH and TLR4ΔBM MASH mice (Fig 3A bottom; B; C). Those mice also had elevated expression of fibrosis-related genes *Col1a1, Lox*, and *Timp1* compared to controls and no differences between TLR4ff MASH and TLR4ΔHep MASH mice (Fig 3D-F).

**Figure 3:**
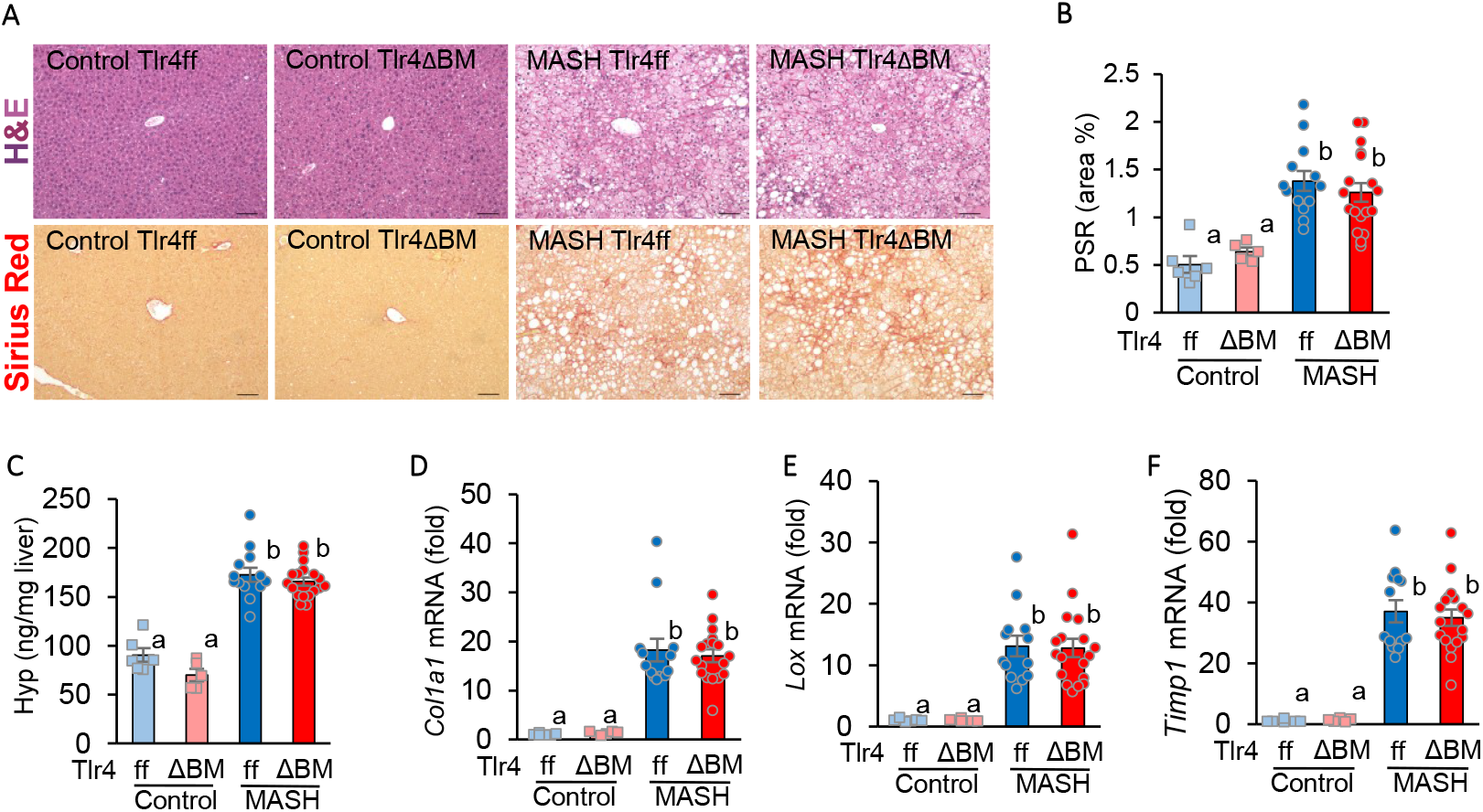
Tlr4 deletion in bone marrow-derived cells does not prevents fibrosis in MASH. Mice with a *Tlr4* deletion in bone marrow-derived cells (TLR4ΔBM) or having a floxed allele and expressing *Tlr4* (ff) were either Ay mice fed a HFFD to induce MASH, or aa fed a low-fat diet to serve as controls. Liver fibrosis was assessed by PSR staining (A, B), hydroxyproline hepatic content (C), and mRNA expression of fibrosis-related genes *Col1a1, Lox*, and *Timp1* (D-F). N = 8/10/15/16, one-way ANOVA; different superscript letters indicate statistically significant differences with p<0.05.

Therefore, these data show that TLR4 signaling in bone-derived inflammatory cells does not mediate the development of fibrosis in MASH.

### TLR4 deletion in hepatocytes did not prevent MASH-related fibrosis

To determine whether TLR4 in hepatocytes promotes the development of fibrosis in MASH, we deleted *Tlr4* in hepatocytes by administering AAV8-TBG-Cre to *Tlr4* floxed mice (TLR4ΔHep). *Tlr4* floxed mice that received AAV8 *null* adenoviruses were included as TLR4-expressing controls (TLR4ff). We included both TLR4ΔHep and TLR4ff Ay mice in which we induced MASH as well as lean control mice.

TLR4ff MASH and TLR4ΔHep MASH mice developed similar levels of obesity and MASH, with comparable liver steatosis, hepatocellular injury, and inflammation (data not shown). TLR4ff MASH and TLR4ΔHep MASH mice both developed fibrosis, as assessed by picro-sirius red collagen staining and hepatic hydroxyproline quantification and compared with controls (Fig 4A-C). Those mice also expressed increased levels of fibrosis-related genes *Col1a1, Lox*, and *Timp1* (Fig 4D-F). However, there was no difference between *Tlr4*-deleted (TLR4ΔHep) mice and *Tlr4* expressing (TLR4ff) mice.

**Figure 4:**
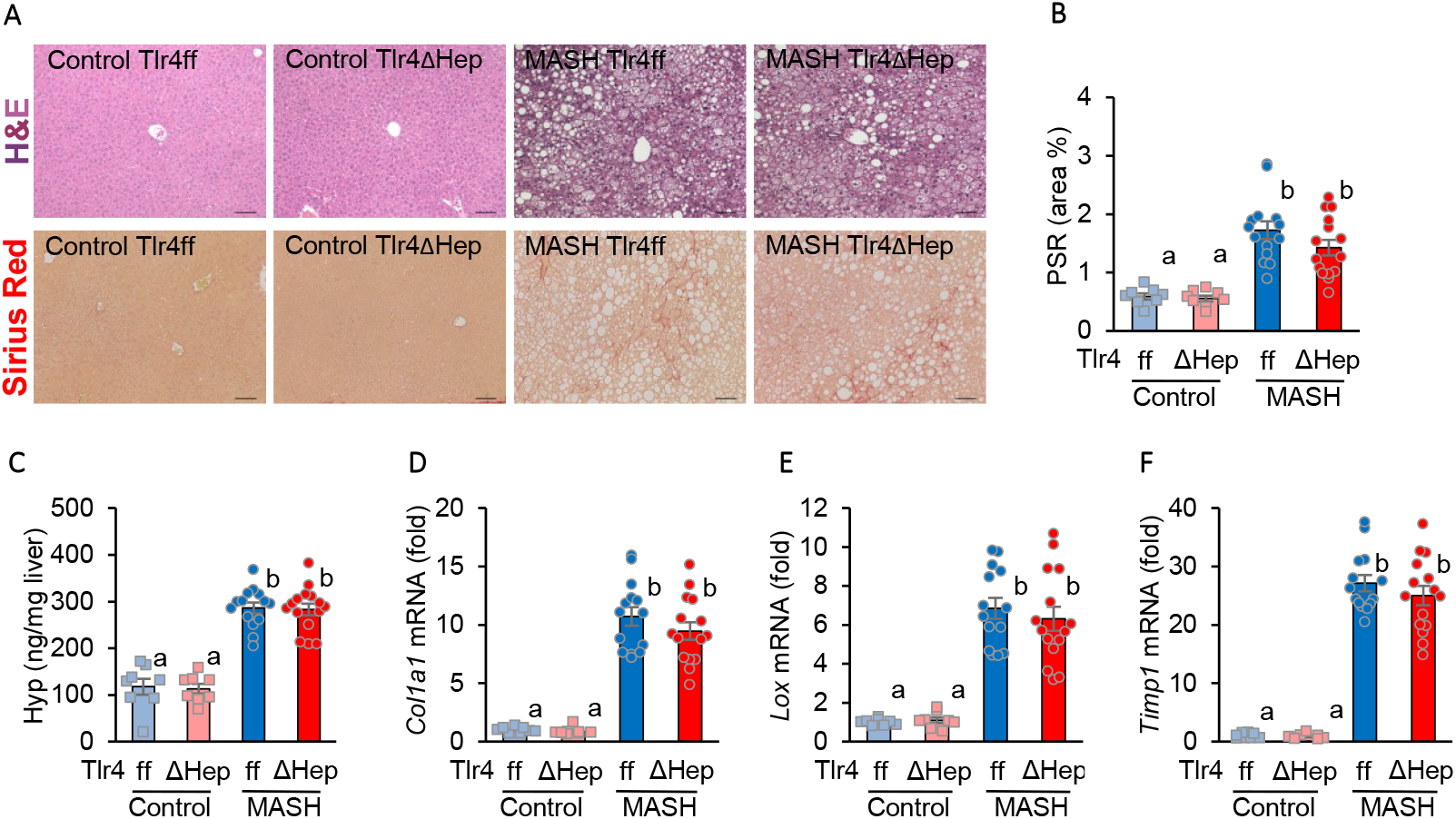
*Tlr4* deletion in hepatocytes does not prevents fibrosis in MASH. Mice with a *Tlr4* deletion in hepatocytes (TLR4ΔHep) or having a floxed allele and expressing *Tlr4* (ff) were either Ay mice fed a HFFD to induce MASH, or aa fed a low-fat diet to serve as controls. Liver fibrosis was assessed by PSR staining (A, B), hydroxyproline hepatic content (C), and mRNA expression of fibrosis-related genes *Col1a1, Lox*, and *Timp1* (D-F). N = 8/8/14/15, one-way ANOVA; different superscript letters indicate statistically significant differences with *p*<0.05.

Therefore, these results indicate that TLR4 in hepatocytes does not have an essential role in the development of fibrosis in MASH.

## DISCUSSION

The aim of these studies was to determine the role of TLR4 in the development of fibrosis in MASH by using mice with either whole-body or tissue-specific deletions of the *Tlr4* gene.

*Tlr4 null* mice, with a whole-body deletion, showed a decrease in MASH-related fibrosis, as assessed histologically, by chemical analysis, and by gene expression, indicating that in MASH, TLR4 is essential for the development of fibrosis. These results agree with previous publications using various models of MASLD and mutant mice lacking functional TRL4, in which fibrosis was decreased [12-14].

Next, we investigated in which hepatic cell type TLR4 may be important to promote MASH fibrosis: *Tlr4* deletion in bone marrow-derived immune cells did not prevent fibrosis, suggesting that TLR4 signaling in those cells does not promote fibrosis. The role of TLR4 in immune cells in MASLD/NAFLD has been investigated before: Jia *et al* showed that *Tlr4* deletion in bone marrow-derived cells did not affect steatosis [23]. Similarly, Giles *et al* showed that, at standard temperature, *Tlr4* deletion in bone marrow-derived cells did not affect the development of MASLD/NAFLD [31]. In these studies, fibrosis was not assessed, perhaps because the mouse models used did not induce significant fibrosis. Yang *et al* reported that chimeric mice with bone marrow from *Tlr4 null mice*, developed fibrosis comparable to controls [32]. Therefore, TLR4 in inflammatory cells does not seem to be responsible for promoting fibrosis in MASH.

Our experiments in which we deleted *Tlr4* in hepatocytes did not show a decrease in fibrosis, implying that TLR4 in hepatocytes does not play an essential role in fibrosis development in MASH. These results conflict with data by Yu *et al*, who reported that *Tlr4* deletion in hepatocytes decreased fibrosis [24]. This discrepancy may be due to the use of different MASLD models: Yu *et al* used a diet containing 2% of cholesterol, a relatively high amount, and cholesterol levels in hepatocytes promote liver fibrosis [24, 33]; therefore, when cholesterol in hepatocytes is high, TLR4 may play a greater role in fibrogenesis. Therefore, TLR4 activation in hepatocytes does not seem to be absolutely essential for development of fibrosis, since in our model of MASLD, its deletion in hepatocytes does not prevent fibrosis.

In the liver, stellate cells and endothelial cells also express *Tlr4* and activation in those cells may promote of fibrosis [11, 16]. Hepatic stellate cells, when activated, transdifferentiate into myofibroblasts, which are the main cell responsible for the deposition of fibrosis in liver [16]. In a model of hepatotoxicity, deletion of *Tlr4* in hepatic stellate cells did not prevent fibrosis; however, the role in MASLD has not been studied [34]. Therefore, the role of TLR4 in hepatic stellate cells and endothelial cells in MASLD remains to be investigated.

Strengths of these studies include the use of a MASLD model that closely replicates the human disease by using a diet similar to the average US diet and hyperphagic mice with increased food intake, as found in human obesity, as well as replicating the histological, microbiological, and gene expression characteristics of the human diseases [26, 27]. The use of loss-of-function genetic models to test the role of TLR4 is an additional strength. One limitation is that this study did not include deletion of *Tlr4* in additional hepatic cell populations.

In conclusion, we show that TLR4 signaling promotes liver fibrosis in MASH by mechanisms that do not involve TLR4 in hepatocytes or immune cells. This suggest that, in MASH, activation of TLR4 in other liver cells, possibly hepatic stellate cells or endothelial cells, or in a combination of cells, promotes fibrosis. TLR4 signaling seems to be important in human diseases since polymorphisms in TLR4 have been reported to affect liver fibrosis development [15, 16]. Further research on TLR4 signaling may lead to new therapeutic approaches to treat MASLD.

## ACKNOWLEDGMENTS

This work was supported by National Institutes of Health grants K22CA178098, R03DK101863, and R15DK131627, PSC-CUNY grant 62551-00 50, a Brooklyn College Student Technology Fees award, and start-up funds from Brooklyn College (to JMC). *Tlr4 null* mice were provided by Ekihiro Seki (UCLA); *Tlr4* floxed mice were provided by Joel Elmquist (UT Southwestern).

## Footnotes

3 Abbreviations: HFFD: high fat and fructose diet; MASH: Metabolic dysfunction-associated steatohepatitis; MASLD: metabolic dysfunction-associated steatotic liver disease; NAFLD: non-alcoholic fatty liver disease; TLR4: toll-like receptor

## REFERENCES

[1] Lee B P, Dodge JL, Terrault NA. National prevalence estimates for steatotic liver disease and subclassifications using consensus nomenclature. Hepatology 2024; 79:666–673.

[2] Younossi Z M, Kalligeros M, Henry L. Epidemiology of Metabolic Dysfunction-Associated Steatotic Liver Disease. Clin Mol Hepatol 2024.

[3] Rinella M E, Neuschwander-Tetri BA, Siddiqui MS, Abdelmalek MF, Caldwell S, Barb D et al. AASLD Practice Guidance on the clinical assessment and management of nonalcoholic fatty liver disease. Hepatology 2023; 77:1797–1835.

[4] Rinella M E, Lazarus JV, Ratziu V, Francque SM, Sanyal AJ, Kanwal F et al. A multisociety Delphi consensus statement on new fatty liver disease nomenclature. Hepatology 2023; 78:1966–1986.

[5] El-Serag H B. Hepatocellular carcinoma. N Engl J Med 2011; 365:1118–1127.

[6] Simon T G, Roelstraete B, Khalili H, Hagström H, Ludvigsson JF. Mortality in biopsy-confirmed nonalcoholic fatty liver disease: results from a nationwide cohort. Gut 2021; 70:1375–1382.

[7] Filliol A, Saito Y, Nair A, Dapito DH, Yu L, Ravichandra A et al. Opposing roles of hepatic stellate cell subpopulations in hepatocarcinogenesis. Nature 2022; 610:356–365.

[8] Pradere J, Troeger JS, Dapito DH, Mencin AA, Schwabe RF. Toll-like receptor 4 and hepatic fibrogenesis. Semin Liver Dis 2010; 30:232–244.

[9] Huebener P, Schwabe RF. Regulation of wound healing and organ fibrosis by toll-like receptors. Biochim Biophys Acta 2013; 1832:1005–1017.

[10] Żeromski J, Kierepa A, Brzezicha B, Kowala-Piaskowska A, Mozer-Lisewska I. Pattern Recognition Receptors: Significance of Expression in the Liver. Arch Immunol Ther Exp (Warsz) 2020; 68:29.

[11] Seki E, De Minicis S, Osterreicher CH, Kluwe J, Osawa Y, Brenner DA et al. TLR4 enhances TGF-beta signaling and hepatic fibrosis. Nat Med 2007; 13:1324–1332.

[12] Csak T, Velayudham A, Hritz I, Petrasek J, Levin I, Lippai D et al. Deficiency in myeloid differentiation factor-2 and toll-like receptor 4 expression attenuates nonalcoholic steatohepatitis and fibrosis in mice. Am J Physiol Gastrointest Liver Physiol 2011; 300:433.

[13] Liu J, Zhuang Z, Bian D, Ma X, Xun Y, Yang W et al. Toll-like receptor-4 signalling in the progression of non-alcoholic fatty liver disease induced by high-fat and high-fructose diet in mice. Clin Exp Pharmacol Physiol 2014; 41:482–488.

[14] Sutter A G, Palanisamy AP, Lench JH, Jessmore AP, Chavin KD. Development of steatohepatitis in Ob/Ob mice is dependent on Toll-like receptor 4. Ann Hepatol 2015; 14:735–743.

[15] Li Y, Chang M, Abar O, Garcia V, Rowland C, Catanese J et al. Multiple variants in toll-like receptor 4 gene modulate risk of liver fibrosis in Caucasians with chronic hepatitis C infection. J Hepatol 2009; 51:750–757.

[16] Guo J, Friedman SL. Toll-like receptor 4 signaling in liver injury and hepatic fibrogenesis. Fibrogenesis Tissue Repair 2010; 3:21.

[17] Cani P D, Amar J, Iglesias MA, Poggi M, Knauf C, Bastelica D et al. Metabolic endotoxemia initiates obesity and insulin resistance. Diabetes 2007; 56:1761–1772.

[18] Ebbert J O, Jensen MD. Fat depots, free fatty acids, and dyslipidemia. Nutrients 2013; 5:498–508.

[19] Holland W L, Bikman BT, Wang L, Yuguang G, Sargent KM, Bulchand S et al. Lipid-induced insulin resistance mediated by the proinflammatory receptor TLR4 requires saturated fatty acid-induced ceramide biosynthesis in mice. J Clin Invest 2011; 121:1858–1870.

[20] Shi H, Kokoeva MV, Inouye K, Tzameli I, Yin H, Flier JS. TLR4 links innate immunity and fatty acid-induced insulin resistance. J Clin Invest 2006; 116:3015–3025.

[21] Machida K, Tsukamoto H, Mkrtchyan H, Duan L, Dynnyk A, Liu HM et al. Toll-like receptor 4 mediates synergism between alcohol and HCV in hepatic oncogenesis involving stem cell marker Nanog. Proc Natl Acad Sci U S A 2009; 106:1548–1553.

[22] Khanmohammadi S, Kuchay MS. Toll-like receptors and metabolic (dysfunction)-associated fatty liver disease. Pharmacol Res 2022; 185:106507.

[23] Jia L, Vianna CR, Fukuda M, Berglund ED, Liu C, Tao C et al. Hepatocyte Toll-like receptor 4 regulates obesity-induced inflammation and insulin resistance. Nat Commun 2014; 5:3878.

[24] Yu J, Zhu C, Wang X, Kim K, Bartolome A, Dongiovanni P et al. Hepatocyte TLR4 triggers inter-hepatocyte Jagged1/Notch signaling to determine NASH-induced fibrosis. Sci Transl Med 2021; 13:eabe1692.

[25] Lorenz E, Mira JP, Frees KL, Schwartz DA. Relevance of mutations in the TLR4 receptor in patients with gram-negative septic shock. Arch Intern Med 2002; 162:1028–1032.

[26] St Rose K, Yan J, Xu F, Williams J, Dweck V, Saxena D et al. Mouse model of NASH that replicates key features of the human disease and progresses to fibrosis stage 3. Hepatol Commun 2022; 6:2676–2688.

[27] Khoj D, Huang R, Altvater E, Ishfaq ZN, Jiang X, Axen KV et al. Mouse Model of Metabolic Dysfunction-Associated Steatotic Liver Disease with Fibrosis. J Vis Exp 2025.

[28] Hoshino K, Takeuchi O, Kawai T, Sanjo H, Ogawa T, Takeda Y et al. Cutting edge: Toll-like receptor 4 (TLR4)-deficient mice are hyporesponsive to lipopolysaccharide: evidence for TLR4 as the Lps gene product. J Immunol 1999; 162:3749–3752.

[29] Schwartz D M, Wolins NE. A simple and rapid method to assay triacylglycerol in cells and tissues. J Lipid Res 2007; 48:2514–2520.

[30] Giles D A, Moreno-Fernandez ME, Stankiewicz TE, Graspeuntner S, Cappelletti M, Wu D et al. Thermoneutral housing exacerbates nonalcoholic fatty liver disease in mice and allows for sex-independent disease modeling. Nat Med 2017; 23:829–838.

[31] Yang L, Miura K, Zhang B, Matsushita H, Yang YM, Liang S et al. TRIF Differentially Regulates Hepatic Steatosis and Inflammation/Fibrosis in Mice. Cell Mol Gastroenterol Hepatol 2017; 3:469–483.

[32] Wang X, Cai B, Yang X, Sonubi OO, Zheng Z, Ramakrishnan R et al. Cholesterol Stabilizes TAZ in Hepatocytes to Promote Experimental Non-alcoholic Steatohepatitis. Cell Metab 2020; 31:969–986.e7.

[33] Ge X, Antoine DJ, Lu Y, Arriazu E, Leung T, Klepper AL et al. High mobility group box-1 (HMGB1) participates in the pathogenesis of alcoholic liver disease (ALD). J Biol Chem 2014; 289:22672–22691.

